# Microbial metabolically cohesive consortia and ecosystem functioning

**DOI:** 10.1101/859421

**Authors:** Alberto Pascual-García, Sebastian Bonhoeffer, Thomas Bell

## Abstract

Recent theory and experiments have reported a reproducible tendency for the coexistence of microbial species under controlled environmental conditions. This observation has been explained in the context of competition for resources and metabolic complementarity given that, in microbial communities, many excreted by-products of metabolism may also be resources. Microbial communities therefore play a key role in promoting their own stability and in shaping the niches of the constituent taxa. We suggest that an intermediate level of organisation between the species and the community level may be pervasive, where tightly-knit metabolic interactions create discrete consortia that are stably maintained. We call these units Metabolically Cohesive Consortia (MeCoCos) and we discuss the environmental context in which we expect their formation, and the ecological and evolutionary consequences of their existence. We argue that the ability to identify MeCoCos would open new avenues to link the species-, community-, and ecosystem-level properties, with consequences for our understanding of microbial ecology and evolution, and an improved ability to predict ecosystem functioning in the wild.

## Community-level metabolism determines broad functions

The importance of microbial communities (MCs)^1^ is increasingly recognised because their control may have enormous benefits for human and environmental health [1–3]. However, our understanding of the complexity of the ecological and evolutionary mechanisms governing the assembly and functioning of MCs is still in its infancy. The notion of MC functioning that we consider here focuses on ecosystem-level processes that operate at temporal scales that are much longer than bacterial generation times. For instance, the carbon cycle in the biosphere happens at geological time-scales, and is contingent on microbial activity [3]. A second feature we consider, is that functioning is not performed by a single species (e.g. a single pathogen) or explained by a single specific process (e.g. formation of a biofilm) [4]. We are interested in broad functions that are the consequence of alternative MC states, which require considering the combined action of multiple taxa, rather than narrow functions, that are the consequence of a single action or metabolic process [5]. Examples of broad functions are human diseases like inflammatory bowel disease or obesity, for which there is evidence that the consequences of MC functions (here health and disease states occurring over the scale of a human lifetime) are unlikely to be explained by a single pathogen but through whole MC states [6].

Similar to studies of macroscopic organisms, the vast majority of microbial studies focus on biomass productivity as a measure of ecosystem functioning [7]. Nevertheless, MCs may vary in their capacity to undertake other functions despite having similar productivities [8]. A common feature of microbial species is their ability to modify the biochemistry of their environment by degrading compounds extracellularly, by secreting antibiotics and signalling molecules, and by uptaking nutrients or releasing byproducts of metabolism. The outcome of MC activity can be defined by the community metabolome and the compounds consumed, which to a great extent can be used to understand how MCs impact broad ecological functions [9, 10].

The increasing availability of whole microbial genomes has boosted the development of detailed mechanistic models of metabolism of single species, for example using Flux Balance Analysis [11]. Recent advances include the incorporation of several species interacting over time [12] and space [13]. With these models it is possible to develop *bottom-up* approximations that predict community-level functioning. Starting with a curated metabolic model of every species in the community (the bottom), it is possible to combine metabolic models to create a larger virtual space in which the metabolites are interchanged, generating a community-level simulation (i.e. upwards) of metabolite fluxes and their associated ecological functions. One difficulty with these approximations is their poor scalability with increasing species number. Although recent methods have enormously simplified the process [14], numerous challenges remain in ensuring the robustness of community-level results against inaccuracies in the single-species metabolic models, and in understanding how historical contingencies and environmental stochasticity during community assembly influence outcomes. While these approximations hold great promise, we are still far from using this technology to get reliable community-level predictions for complex communities. However, we agree with the central premise that metabolism holds the key to linking ecological dynamics (the abundance of species) and ecosystem functioning.

## Complexity, stability, and functioning of microbial communities

An alternative approach arises from the classical problem in theoretical ecology of how the number of species, their identity and their interactions (i.e. community complexity) are related to their stable coexistence [15]. We first assume that two independent replicates of the same community inhabiting the same conditions will lead to the same levels of ecosystem functioning. This would mean that we could, in principal, achieve a predictable level of functioning by re-creating a MC and its associated environmental conditions. However, slight alterations to the relative abundances, to the environmental conditions, or to the ecological histories could lead to different levels of functioning. In addition, the function of interest (e.g. a disease) is not necessarily a consequence of a unique combination of species: many combinations of species may lead to the same observed function (the disease). These observations imply there is a need to expand the classical complexity-stability problem to encompass functioning, in which the focus is moved towards the stability of a function rather than solely the stability of the community.

The likelihood that two distinct MCs lead to the same function depends on the amount of functional redundancy among species. There have been three views, which are not mutually exclusive. The first is that redundancy is widespread due to the prevalence of neutral or convergent evolution, which could lead to functionally redundant species, and therefore the removal of some species will be compensated by others [16, 17]. The second is that the apparent widespread redundancy of rare species [18, 19] is simply due to an inability to measure a sufficient number of community-level quantities to fully characterise their function [20]. A third intermediate view argues for the existence of functional groups, whose species members are functionally redundant, but that combining members of different groups, would lead to different MC functions.

Large-scale patterns may help to differentiate among these scenarios, for instance through the analysis of diversity patterns across spatial or environmental gradients [21]. For example, according to Hubbell’s neutral theory [22], the *β*-diversity should increase across sites with increasing spatial distance if there is limited dispersal whereas there should be no change with respect to environmental gradients since species are functionally equivalent. Conversely, some species may consistently coexist due to similar environmental preferences, or due to mutualisms [23], in which case changes in *β*-diversity would be primarily observed across environmental gradients. Despite the relative success of neutral theory to explain some MC patterns [16], a growing body of research has found persistent MC states likely driven by environmental conditions, including for example the controversial discovery of human enterotypes [24,25]. As an example, we illustrate in Fig. 1A the clustering of more than 700 communities sampled along a spatial gradient spanning 5 orders of magnitude, from 5m to 150km, into just 6 community classes. Ref. [8] showed that there was significant spatial autocorrelation, implying an important role of stochastic processes, see Fig. 1B. Nonetheless, communities joined in these classes were often distant in space, suggesting that *β*-diversity similarity likely driven by local environmental conditions, such as rain and drought. Importantly, when a variety of ecological functions were quantified (respiration, ATP production or degradation of extracellular substrates), the community classes were associated with distinct functional profiles. The classes also had differentiated genetic repertoires, pointing towards a relation between ecological conditions and bacterial traits, and rejecting the hypothesis that the communities were functionally equivalent. Comparable large-scale efforts, including Tara Ocean [26] and the Human Microbiome Project [27], have also found that for natural communities the number of possible states are constrained.

**Figure 1:**
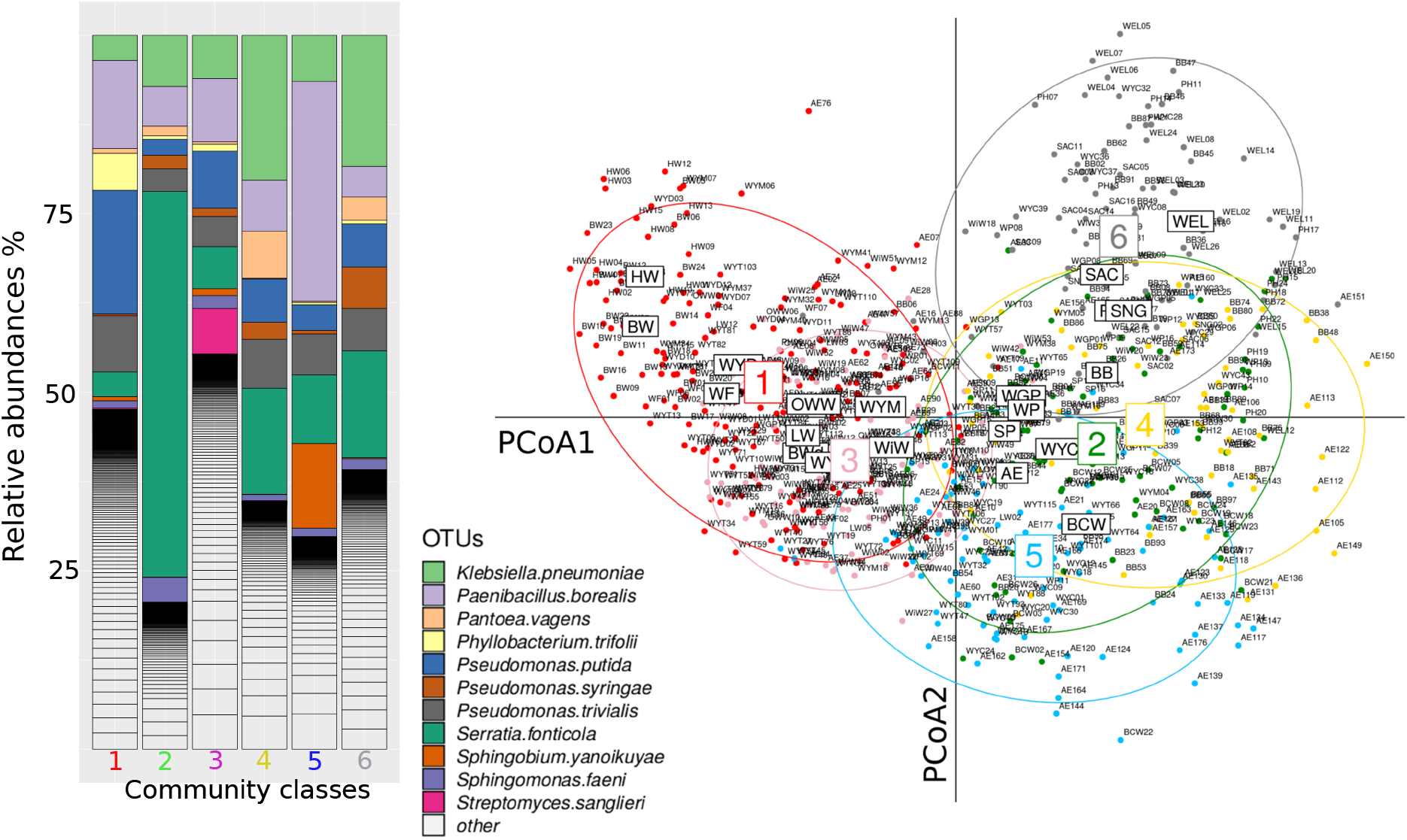
Community-level classes. (Left) Approximately 700 natural MCs sampled from tree-holes can be classified into six classes (left bar-plots). The coloured surfaces of each bar represent the relative abundance of the most representative species (OTUs at >97% sequence similarity), being the total bar a 100%. (Right) Projection of the similarity of the communities into the first two principal coordinates of a Principal Coordinate Analysis. Significant spatial autocorrelation was found, which becomes apparent from the centroids of the locations were the communities were sampled (three letters boxes). However, the classes (labelled 1 to 6 and highlighted with elipsoids) bring a more economical classification with comparable significance, with their communities often sampled from different locations. In addition, communities classified in these classes have distinct functions and metagenomic repertoires, suggesting that local environmental conditions occuring in different locations rather than neutral evolution with dispersal limitation shape these communities. More details in Ref. [8].

Once broad community classes have been identified, the next step is to predict a MC metabolome from the information provided by its metagenome [28–30], and then to step-down into the contributions of constituent species have to predicted metabolome, and therefore their role in the function [29]. Nevertheless, to further explore the relation between complexity, stability and function, approximations that can predict unobserved MCs states from observed states are needed. Despite the apparent existence of simple assembly rules [31, 32], it is difficult to reliably parameterizing multispecies population dynamics models to make predictions even if long time-series samples are available, [33,34], although recent progresses can be found [35, 36]. Since, microbial communities are typically monitored by community sequencing of the 16S rRNA locus, it would be helpful to develop improved methods to parameterize population dynamics models in order to generate predictions of ecological functions.

## Environments can be defined by their resource content

We aim to understand microbial population dynamics, and link those dynamics to functioning via the metabolisms of the community members. One avenue is to classify environments according to the qualitative properties of the resources. We suggest there are three important axes: i) The effective number of different resources, i.e. the heterogeneity of the environment. “Effective” is used here to distinguish resources in terms of their molecular content. For instance, an environment containing chitin and cellulose, would have a lower effective number than chitin and phosphoric acid because the former pair has a more similar composition. ii) The resource abundance iii) The stage of degradation, particularly the compounds, degradation products, and energy that can be retrieved from each unit of effective resource.

To illustrate these axis, in Fig. 2 we show the population dynamics of a marine natural bacterial community colonising synthetic particles of alginate [37]. In the figure, the Exact Sequence Variants (ESV) [38] belonging to the 20 most abundant genera (Operational Taxonomic Units (OTUs) with sequence identity > 95%) are highlighted, with the remaining genera shown in dark grey. The first remarkable feature of this experiment is the high reproducibility of the trajectories (three independent replicates per time point) despite being colonised by a natural community, which we would expect to have a somewhat stochastic assembly. At the starting stages of the assembly of the particles, a few OTUs dominate the community, possibly specialised on the breakdown of alginate. At an intermediate stage, around 60-84h, there is a clear transition in the communities, likely mediated by the release of by-products that are now acquired by secondary consumers. As a consequence, there is an increase in the number of microniches and a consistent increase in the diversity of the community.

**Figure 2:**
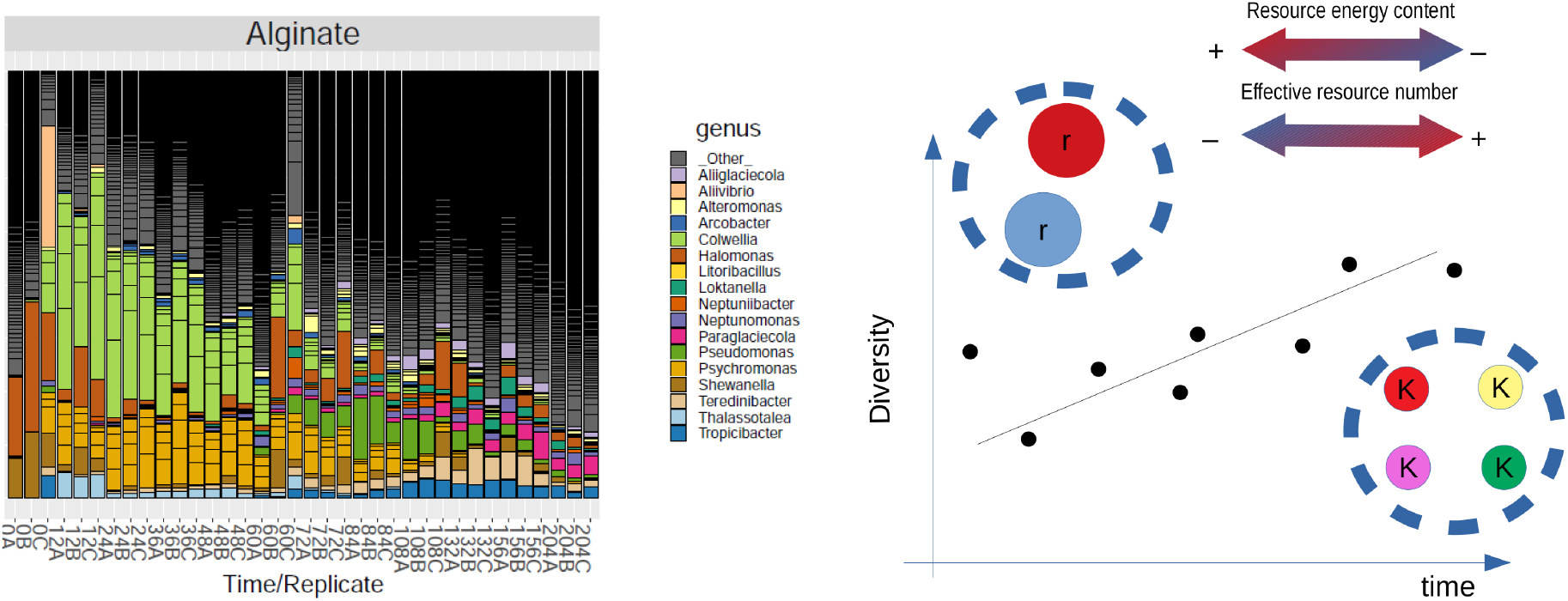
Diversity and resources properties in an ecological succession. Population dynamics of natural marine assemblages MC on synthetic particles of alginate (left). Each bar represents the relative abundance in the correspondening sample, labelled by replicate and time-point. (Right) Illustration of the expected diversity increase through time as a function of the energy content and the effective number of resources that results from the degradation of resources. r-strategists will be observed at earlier times where the resources are abundant and rich in energy, while K-strategists should be expected at later times, where the resources are more heterogeneous, lower in energy and scarce.

This example also illustrates that the three environmental axes we propose are not fully independent. We observe that, as soon as the single abundant polymer present in the beginning is degraded, new lower-energy resources are created, increasing their effective number, consistent with recent theoretical results [39]. Therefore, we expect communities would display a single strategy when the environment remains approximately constant in these three axes, for instance r-strategists in an environment rich in a single complex nutrient, where density-dependent processes are not relevant [40, 41]. On the other hand, we expect K-strategists to dominate a high diversity regime where density-dependent processes such as competition for resources drive the ecological dynamics. And we should expect a scenario of ecological succession otherwise, with different strategies dominating particular time-windows.

## Stability mediated by niche shaping: Metabolically Cohesive Consortia

The qualitative picture presented brings a broad framework to understand community-level succession, but the high predictability during community assembly, as shown in Fig. 2, requires more explanation. To explain this observation, we follow Tilman’s seminal results using consumer-resources models, which predict that the outcome of two species competition for a single resource will be determined by the ability of each species to deplete the resource when grown in isolation. When both species are grown in co-culture, it is expected that the one depleting the resource to a minimum in isolation, will exclude the other in co-culture [42]. As we have discussed above, this scenario rarely occurs in microbial communities because the “winner” releases metabolic by-products that can be used by the “losing” species, which provides the opportunity for coexistence if the outcompeted species is able to exploit this new niche [43, 44]. This reasoning can be extended to a large number of species and resources. We suggest that one hypothesis that emerges from this observation is that the combination of species that maximally deplete resources to a minimum will systematically dominate the community. A corollary is that such community will optimize functions associated with the community-level metabolism, as recently shown for methanogenic communities [45].

This dynamical formation of reproducible modules is supported by several theoretical results working with consumer-resource models, which have reported the systematic formation of stable communities starting from random assemblages [39,46–49], via resources partitioning, and even when the active release of by-products is absent [50].

To illustrate this point, we discuss a recent study investigating a consortium commonly found in the production of kefir and wine formed by *Sacharomyces cerevisiae* and two *Lactobacillus* species (LAB: *L. lactis* and *L. plantarum*). The two LAB species are auxotrophs of some amino-acids that are provided by *S. cerevisiae* in nitrogen-rich environments [51]. In addition, when the carbon source is lactose, the inability of *S. cerevisiae* to grow on this carbon source is compensated by a supply of glucose from *L. lactis* [51]. Therefore, a mutualistic relationship between these species makes the consortium robust against fluctuations in the available amino-acids or carbon sources in the environment. This occurs through a niche created from overflown metabolism [51], which may rapidly result in a stable consortium. Therefore, niche self-construction would be an expected ecological consequence of the collective reduction of environmental (i.e. resource) fluctuations.

We call this kind of community organisation a **Me**tabolically **Co**hesive **Co**nsortium (MeCoCo), which is a type of consortium that exhibits a positive feedback loop, where the consortium engineers the environment by altering resource abundances, and stabilise its dynamics. Members of MeCoCos are specialised as a whole, and effectively minimise competition within the consortium and exclude resource generalists by lowering resource abundances through their combined action. We conjecture that the formation of MeCoCos may be common in natural communities, and would be a parsimonious explanation for the observation of clustered community compositions at broad scales (Fig. 1), and the predictable successional trajectories exhibited in Fig. 2.

Still, the reproducibility of the observations require taxa-specific outcomes. While in most theoretical models stable assemblages result from combinations of species with random metabolic strategies, in reality microbial species have specific capabilities, which would lead to the observed specificity in terms of taxonomical composition via co-selection of species that jointly optimize the depletion of resources of the consortium [45]. The time-windows in which evolutionary and ecological proccesses happen may also be relevant for the formation of MeCoCos. For instance, a systematic formation of modules with reproducible composition is a result predicted for low immigration rates [48]. Moreover, it has been recently shown with simulations that stable consortia often emerge in complex communities containing large population sizes or high mutation rates [52], in which evolutionary events in the population may occur faster than the time that the population has to equilibrate the ecological dynamics. These consortia can then spatially spread, outcompeting resident communities. Therefore, the ecological dynamics has a strong influence on the environment in the MeCoCo hypothesis. For large population sizes or low immigration rates we may expect that species rapidly coevolve towards the emergence of efficient consortia. This possibility would relax the necessity of finding the specific environmental requirements or a specific combination of species since we would expect MeCoCos to be the outcome of complex communities competing for resources that are gradually degraded.

## Empirical evidence of MeCoCos

There is a large body of literature reporting the stable coexistence of species pairs through cross-feeding, with metabolic and physiological mechanisms thoroughly described in [53, 54] and [55] respectively, and hence will not be discussed here. *In vitro* experiments with larger consortia can be found with genetically modified strains such as in [13], and through the generation of auxotroph strains [56]. It is challenging to create synthetic assemblages of more complex communities because it is difficult to generate the appropriate environmental conditions that mimic those encountered in natural environments [57]. Finally, if a MeCoCo exists in nature, it is challenging to capture the whole MeCoCo using exclusively culture-based approaches, and the absence of a small number of species might prevent the proper functioning of the consortium as a whole [58].

Aside from culture-based approaches, there is growing evidences of the existence of large consortia in natural environments. Although there are many results in controlled biotechnological settings or conditions such as anaerobic reactors [45], generation of biofuel [59] or pesticides degradation [60], a more comprehensive ecological framework is needed to understand complex processes in natural environments, including for example global biogeochemical cycles. Examples of this kind are consortia of strains from at least 18 genera that inhabit dental plaques [61], consortia between phototrophic green sulfur bacteria and chemotrophic bacteria such as *Chlorochromatium aggregatum* [62], and anaerobic food chains in methanogenic environments [63].

We illustrate the prominence of MeCoCos by expanding the alginate-particle experiment shown in Fig. 2 to also include particles made of agarose or a mixture of agarose and alginate (Fig. 3) (Ref. [37]). We observed highly reproducible dynamics in all three conditions, with a shift in the composition at intermediate times, and partitioning of the time-series into two or three groups of distinct species likely due to an abrupt transition in the underlying substrate. Despite being different substrates, this consistent dynamic occurred in all three experiments but with different community members, suggesting a tight relationship between the substrate and the communities. Moreover, the abundances of the most representative species in particles composed of a mix of alginate and agarose (Fig. 3, right) experienced dynamics that could be approximated by a simple linear combination of their dynamics in the single substrates particles (Fig. 3 left and middle) [37]. We note that the communities occupying the particles were composed of just a few genera, which would be our candidates for MeCoCos in this dynamical environment, with most of the genera involved in more degraded substrates.

**Figure 3:**
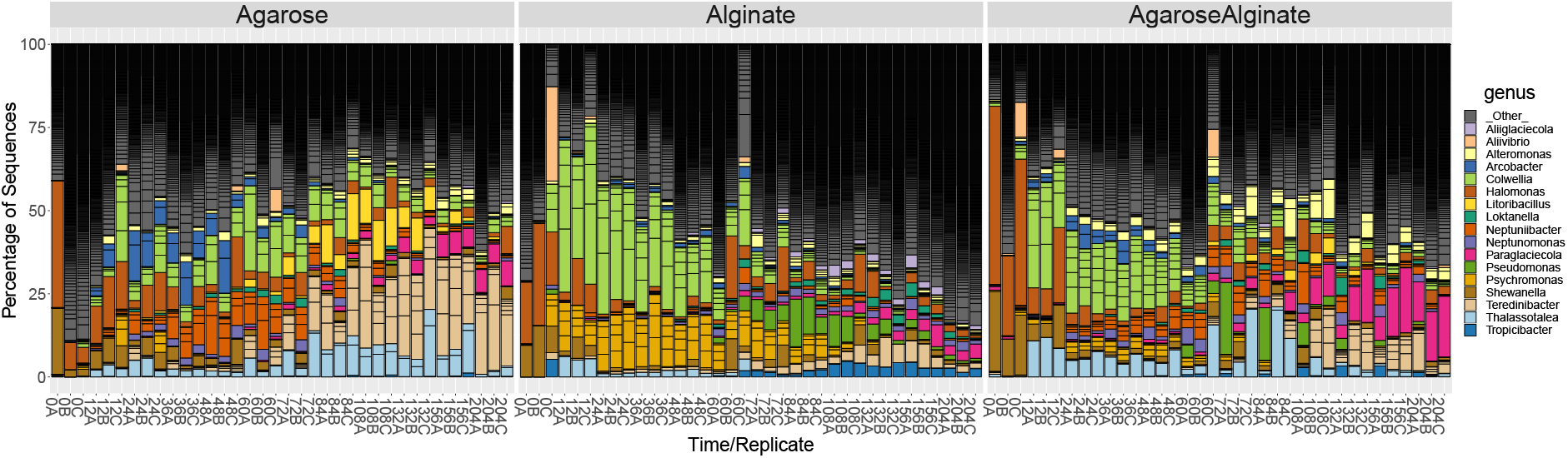
Population dynamics in single- and mixed- resource environments. Microbial communities were allowed to colonise particles composed of alginate (left), agarose (middle), or a mixture of two (right). The community dynamics in all three conditions exhibit a transition at intermediate times, in which the dynamics is split into well diferentiated communities. Although the community composition on the two substrates difers, the remarkable modularity of the experiments allows us to predict the relative abundance for the most abundant members as a linear combination of the relative abundances of the pure substrates (Ref. [37]).

In examples including the dental plaque ecosystem, it is necessary for the community members to be in close proximity, even in direct contact [55]. Therefore, an open the question is whether syntrophy can be maintained in more dynamic or open environments [64]. Consistent regularities in community composition observed in the analysis of large datasets of natural samples interrogated with 16S rRNA sequencing, provide some support to the existence of MeCoCos in many habitats. In controlled environments, experiments analysing the growth of natural communities reported a systematic convergence to 3 the same steady states at the family level [65].

## Ecological consequences of MeCoCos

An important corollary to the MeCoCo hypothesis is the prediction that these consortia are stable against invasions because no other species would be able to uptake the necessary nutrients at a suffciently high rate to coexist with the established members. There is a growing body of results in this direction, highlighting the importance of metabolic interactions to shape stable communities [69–71].

If MeCoCos are stable structures that resist invasion, it would be possible to understand microbial ecosystems in terms of MeCoCo building blocks. To illustrate this point, we outline a hypothetical example in which we consider groups of species with similar metabolic capabilities (see Fig. 4, left), which we call functional groups because this is the definition of function that we have adopted here. In this example, we would typically expect competitive interactions within these functional groups (represented in the networks with dotted lines) because members occupy similar metabolic niches [72]. Similarly, we may expect that metabolites are traded between difierent functional groups, leading to commensal or mutualistic interactions (continuous lines in the network). With this picture in mind, we may think that competing species tend to segregate while those cooperating tend to aggregate. This would allow us to rearrange the network to observe that a new level organisation becomes apparent (see Fig. 4, right). In this visualisation, there is intra-specific cooperation within MeCoCos, while competition occurs between MeCoCos. Interestingly, although we believe that metabolic interactions are critical, other kind of interactions may also be accommodated. For instance, bacteria may also be organised into social cooperation due to their collective antibiotic resistance, while antibiotic-mediated antagonism may occur between populations [73].

**Figure 4:**
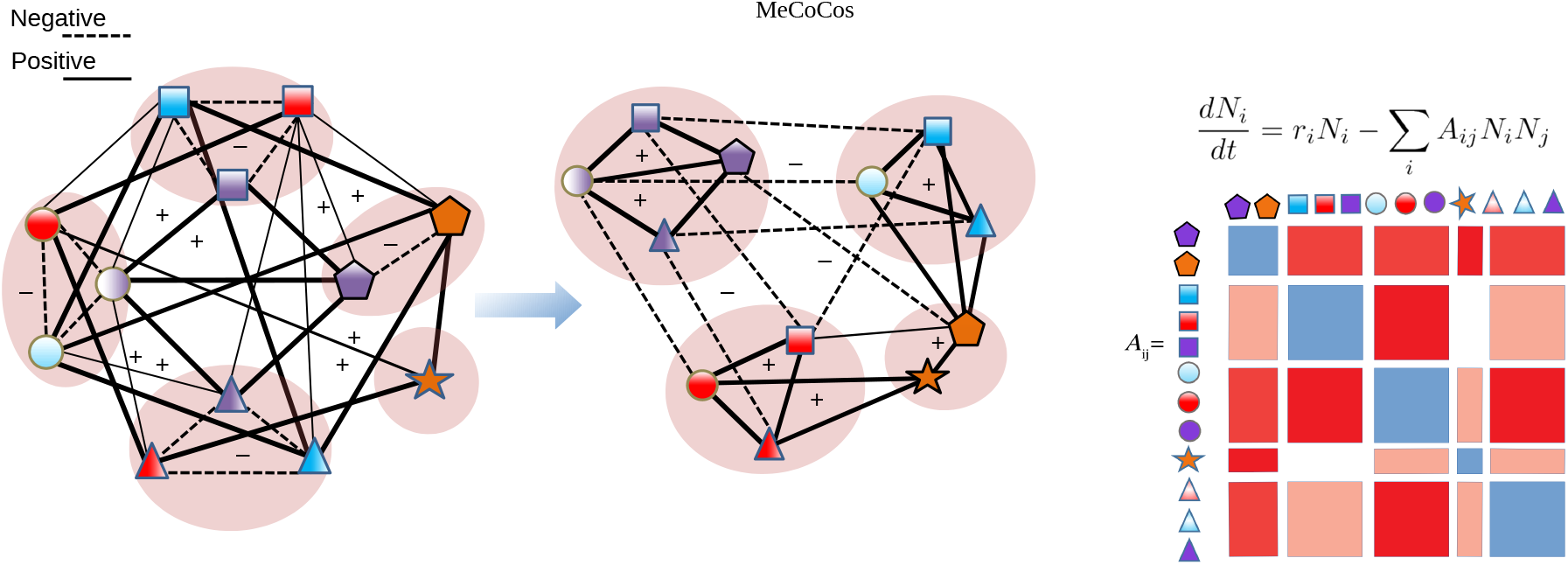
MeCoCos as a new effective level of organisation. (Left) The symbols represent different species, where the same shape represents similar metabolic capabilities (left network, functionally redundant groups), which would lead to competitive interaction (dotted lines) due to a high niche overlap. On the other hand, members of different functional groups may engage in commensal or mutualistic relationships, driven by metabolic complementarity (continuous lines). A rearrangement of the left network leads to a new representation (right network) in which members related through complementary functions tend to co-occur together, forming MeCoCos, which constitute a new intermediate level of organisation between the species- and the community-levels. Understanding community-level dynamics may thus be simplified to understanding how MeCoCos effectively compete (see Fig. 4). (Right) The interaction matrix that results from these networks has a block structure that makes it suitable to build effective population dynamics models as in macroscopic systems (e.g. [66–68]) Blue blocks represent competitive interactions, and red blocks represent mutualistic interactions arising from metabolic complementarity.

The MeCoCo that prevails in competition with other MeCoCos will depend on the environmental conditions, but we would need to understand the behaviour of MeCoCos as a whole, and not of every single species. Also the specific members ofMeCoCos are likely context-dependent [6] with relationships between particular species more likely found than others. In particular, rare species might be seen as “metabolic switches” allowing for the formation of specialised metabolic pathways under certain conditions (Fig. 5).

**Figure 5:**
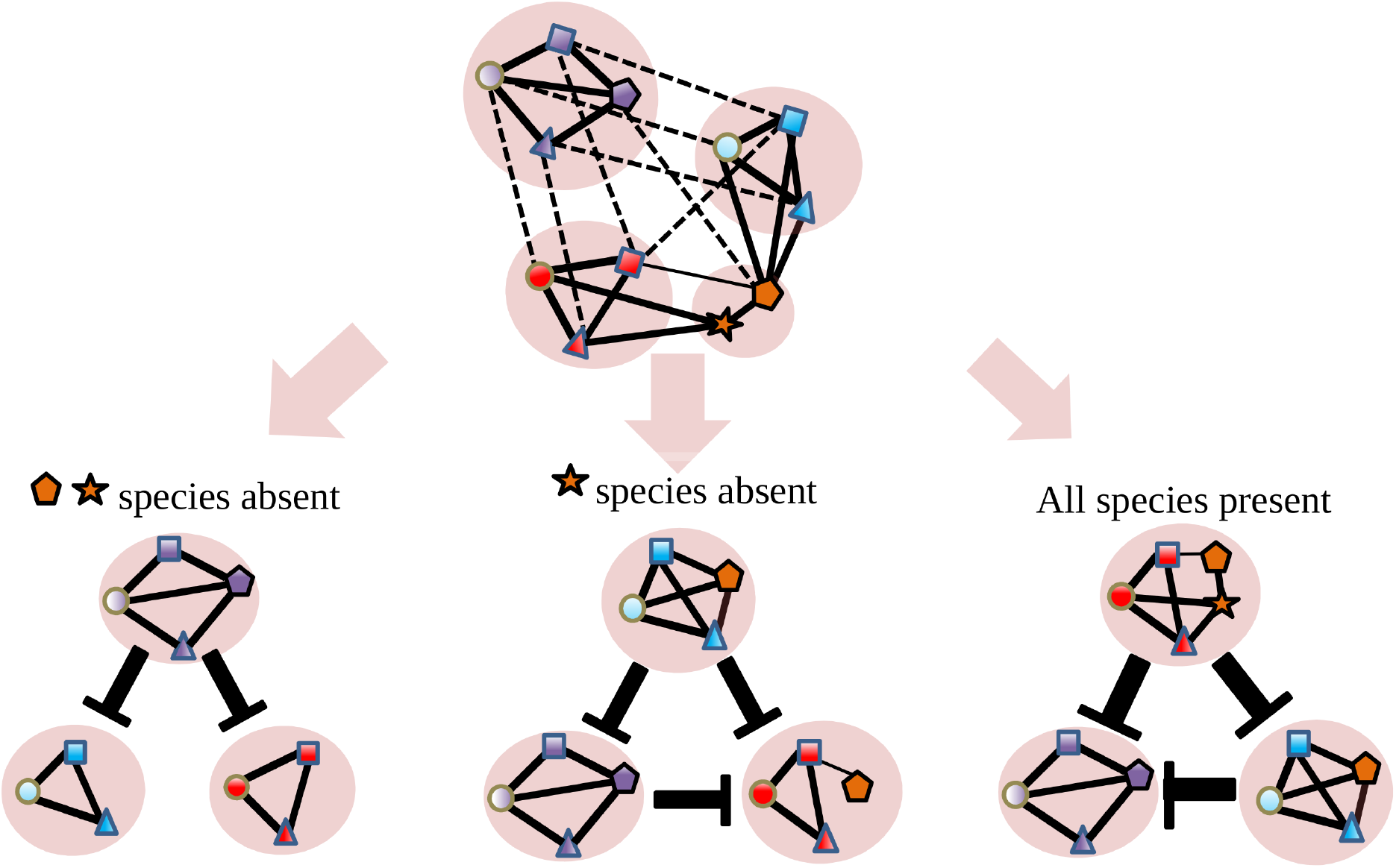
Competition between MeCoCos. Illustration of how changes in species composition alters which MeCoCo becomes dominant. The dominant community is the one that depletes resources to a minimum, which we assume is related with the number of realised metabolic (complementary) links (continuous lines connecting species). The orange species may be understood as community-level “metabolic switches” that determine the outcome of the dynamics.

Studying MeCoCo building blocks in this manner would facilitate the modelling the outcome of community coalescent events, when two microbial communities are mixed together [74]. Typical examples of these events are the mix of oral communities when kissing, or a tree transplanted into new soil. There is growing interest in exploiting these community-mixing events, most notably in recent years the transplant of human faeces received much attention [75].

In general, the framework proposed would move the emphasis from species to MeCoCos, and in so doing would reduce the complexity in predicting the outcome of coalescent events, allowing coalescense to be modelled with population dynamics models at the MeCoCos level. In Fig. 4 we show how the MeCoCo structure leads, in terms of an interaction matrix, to a block structure similar to the one found for trophic or mutualistic macroscopic systems. This would allow us to simplify these systems to reasonable levels of complexity, in the same manner as for macroscopic systems [66], for example dividing ecosystems into mutualistic interactions between plants and their pollinators and competitive interactions within plant and pollinator functional groups [67, 68].

## Evolutionary consequences of MeCoCos

The large number of metabolic genes found in the prokaryotic flexible genome is consistent with a picture in which metabolic trading is widespread in natural environments [76,77]. Under the MeCoCo hypothesis, the genomic regions that are critical for the maintenance of the species in the consortium will be subjected to a strong selective pressure. Conversely, those genes associated with functions that are covered by other species will experience reduced selective pressures, akin to that observed in symbiotic or parasitic species, leading to a rapid reduction in genome size. The role of selective pressures in structuring MeCoCos could be governed by effective population sizes. Low *N*_*e*_ would favour genetic drift and hence the appearance of genomic structures that cannot be purged by selection, like pseudogenes. If we consider that the formation of MeCoCos in natural environments is frequent, we hypothesise that their members have large *N*_*e*_, in which case the selection pressure is larger and the loss of genes should have a more immediate selective advantage.

Another scenario in which widespread loss of genes has been observed is in oligotrophic environments like the open ocean, where streamlining theory explain this loss pointing towards the necessity of these species being efficient consumers in environments with low nutrient availability [58]. Under these conditions, there is a strong pressure to maintain an efficient core metabolism and to lose unnecessary genes [58].

The genetic signatures expected for MeCoCos lie in between the scenarios observed for parasites and symbionts, and oligotrophic species. This is the prediction made by a related hypothesis, the Black Queen Hypothesis (BQH), which hypothesises this loss on the basis of the existence of leaky metabolic products frequently present in MCs [78]. For the BQH hypothesis, the loss of genes may be seen as a passive consequence of the fact that these products (e.g. metabolites) are public goods, leading to the loss of the metabolic pathways needed for their biosynthesis in taxa that can uptake the metabolites from the environment.

In the case of MeCoCos, we expect a more dynamic eco-evolutionary process in which there is selection not only to lose genes that are redundant within the MeCoCo, but also to promote genes that maintain the consortium. Therefore, we would expect both auxotroph species, and also the selection of genes related with other functions that may lie beyond metabolism, such as chemotaxis and signalling to identify partner species and to stay in close proximity [55].

## Conclusion

In this article we discussed different approaches to understand community-level function in microbial communities. We argue that there is a need to focus on the relationship between the complexity of the community (the number and identity of the species and their interactions) and community-level metabolism. However, instead of approaching this relationship from the bottom-up by building detailed models for each member species, we advocate for top-down approaches, in which natural communities are isolated and manipulated under controlled conditions.

The rationale behind this choice is that bottom-up approaches require accurate characterisation of every species’ metabolism and to develop strategies to tackle the combinatorial explosion arising from all possible ways in which species can be combined to build a community. Top-down approaches aim to learn instead from natural patterns, and then to step down into the mechanistic processes.

In this respect, an important property to look at is the stability of the communities over space and time, under the assumption that stable communities will lead to functions that are also stable in time. Nevertheless, we pointed out that the relationship between the complexity of a community and its metabolome is many-to-one, so it is important to explore patterns of functional redundancy in communities. To address this task we think that is not only important to increase the number and resolution of functional measurements, but to move away from a descriptive characterisation and i) perform experiments on which we can monitor function while manipulating the ecosystem [79, 80]; and ii) we need a more mechanistic characterisation of MC functions in these experiments, as in [70, 81].

From these kinds of experiments, we highlighted the necessity of finding a broad ecological and evolutionary framework in which a more mechanistic understanding of the relationship between diversity and function can be developed. Given the ability of bacterial communities to modify the environments they inhabit, we postulated that bacteria may commonly construct Metabolically Cohesive Consortia (MeCoCos) through complementary syntrophic interactions. One outcome of this relationship is the increased control of the existing resources by members of the consortium, which would promote the stability of the community and of functioning. We predict that MeCoCos would deplete resources to a minimum and would be more stable against invasions. In addition, MeCoCos should exhibit high levels of structural stability, such that fluctuations in external input of resources would be buffered by species producing the resource or its derivatives, a type of stability whose importance has been emphasized in macroscopic organisms in the context of mutualistic systems [67, 68, 82], and just recently for microbial populations [39, 49, 83]. It is less clear how stability against large unexpected environmental fluctuations would impact other components of stability, such as the persistence of component species [84]. We believe there is great benefit in understanding this new level of organization, which would lie in between the species and the community-level, facilitating the establishment of new bridges between bottom-up and top-down approximations.

Several questions remain, particularly around the identification of MeCoCos in nature. We predict that species belonging to the same MeCoCo should systematically co-occur since certain combinations may be the most successful in shaping their own niche [70], potentially providing a straightforward route to identifying MeCoCos. However, whether this could be widely applied remains to be verified. We also have an incomplete picture of or which are the most likely environmental conditions where we expect MeCoCos to occur, though MeCoCos might be difficult to maintain in the face of high disturbance frequency or high levels of immigration.

If MeCoCos are widespread, there would be a need to integrate this idea into the broader evolutionary literature that discusses levels of selection. If species and their interactions can be understood as portions of metabolic processes that together complete more complex metabolisms [85], metaphors have been used of bacterial supra-organisms [86]. There is likely little need to re-open the debate around the existence of group selection [87, 88] since the formation and maintenance of MeCoCos could be explained following a process based on eco-evolutionary feedbacks [52]. However, there is the possibility to integrate this idea with emerging views of the evolution of multicellularity, since the formation of stable consortiums have been suggested as an intermediate state for the formation of multicellular life [62, 89, 90].

This study was motivated by the need to simplify the extraordinary complexity of microbial life that has been discovered in natural systems so that we can predict and control ecosystem functioning. The idea that much microbial life lives off metabolic biproducts produced by other microbes is as old as modern microbiology, including some of the earliest examples of how microbial ecosystems operate Winowgradski columns. We believe a refinement of this view is warranted-that functions are ultimately driven by self-selecting consortia, driven by the way in which microbes exploit and modify their surrounding environment. Identification of MeCoCos using the wealth of genomic data currently being generated would pave the way for both a simpler and more mechanistic understanding of the link between microbial dynamics and the functioning of ecosystems.

## Methods

Data were retrieved and processed following [5] (Fig. 1) and [91] (Figs. 2 and 3). Bar-plots were generated rarefying samples to 1000 reads and represented with the R package Phyloseq [92, 93]. In Fig. 1 the classes, *β*-diversity distance matrix and GPS locations of the samples were retrieved from [8], and we performed a Principal Coordinate Analysis (PCoA) of the samples with the R function dudi.pco, from package ade4 [94]. Results were represented projecting the samples into the first two PCoA coordinates and computing the centroids of the clusters defined by both the sampling sites and the community-classes.

## Acknowledgements

The research was funded by a European Research Council starting grant (311399-Redundancy) awarded to T.B. and by the Simons Collaboration: Principles of Microbial Ecosystems (PriME), award number 542381, to S.B. T.B. was also funded by a Royal Society University Research Fellowship.

## Footnotes

1 For simplicity, in this article we use the term “bacteria” referring to members of bacteria and archaea kingdoms, while “microbial” may also include eukarya.

